# Female fruit flies use social cues to make egg clustering decisions

**DOI:** 10.1101/2024.07.03.600353

**Authors:** Emily R. Churchill, Emily K. Fowler, Lucy A. Friend, Marco Archetti, Douglas W. Yu, Andrew F. G. Bourke, Tracey Chapman, Amanda Bretman

## Abstract

**Background:** The ability to respond plastically to environmental variation is a key determinant of fitness. Females may use cues to strategically place their eggs, for example adjusting the number or location of eggs according to whether other females are present, driving the dynamics of local competition or cooperation. The expression of plasticity in egg laying patterns within individual patches, i.e., in contact clusters or not, represents an additional, under-researched and potentially important opportunity for fitness gains. Clustered eggs might benefit from increased protection or defence, and clustering could facilitate cooperative feeding. However, increased clustering is also expected to increase the risk of over-exploitation through direct competition. These potential benefits and costs likely covary with the number of individuals present, hence egg clustering behaviour within resource patches should be socially responsive. We investigate this new topic using the fruit fly *Drosophila melanogaster*.

**Results:** Our mathematical model, parameterised by data, verified that females cluster their eggs non-randomly, and increase clustering as group size increases. We also showed that, as the density of adult females increased, females laid more eggs, laid them faster, and laid more eggs in clusters. Females also preferred to place eggs within existing clusters. Most egg clusters were of mixed maternity.

**Conclusions:** Collectively, the results reveal that females actively express plasticity in egg clustering according to social environment cues and prefer to lay in clusters of mixed maternity, despite the potential for increased competition. These findings are consistent with egg clustering plasticity being selected due to public goods-related benefits.

## Background

In natural contexts, social environments and availability of resources are highly variable, exposing populations to fluctuating competition. In response, phenotypic plasticity can play a key role in ensuring individuals maximise their lifetime reproductive success (1, 2). Because optimal strategies may vary with fluctuating levels of competition, individuals that respond to the prevailing environment should show increased fitness. Plasticity in investment decisions should be particularly important for females, who can generally exert greater control over reproductive investment in progeny. For example, females can sometimes control paternity (3–5), the number and sex ratio of offspring via ejaculate manipulation (6–8), and the choice of nesting/oviposition site locations. For females that deposit eggs upon a substrate, egg site location can be a vital determinant of hatching success. Consistent with this, female aggregation and oviposition behaviours of many species appear to be plastic in response to environmental variation (9–12).

### Plasticity in aggregation and egg laying - competition, cooperation, and cheating among females

At increasing population densities, there is expected to be a greater risk of direct competition but also increased potential for cooperation, and the relative importance of the two is expected to differ as individuals progress through life stages. For example, when food is limiting and competition high, eggs and larvae are typically more vulnerable to cannibalism than are adults, due to their more restricted mobility (13–15). On the other hand, in some species, there may be benefits of joining with conspecifics when foraging or hunting. For example, Eurasian Coots (*Fulica atra*) copy each other’s behaviours to reduce foraging costs (16), and larvae in metamorphosing species can benefit from increased efficiencies resulting from communal feeding (17, 18). Females can use the presence of conspecifics as a cue to modify reproductive investment in response to changing social conditions. For example, increasing adult female density reduces offspring production and body size in Western Mosquitofish (*Gambusia affinis*) (19). In contrast, in the Willow Leaf beetle (*Plagiodera versicolora*) fitness effects of increasing adult female density can vary with development, resulting in a decrease in egg hatching, but an increase in egg clutch number and duration of laying period (20).

Among females of species that deposit their eggs upon substrates, it is also often found that eggs are laid in conspecific clusters, e.g. in fish (21), birds (22), reptiles, amphibians (23), and invertebrates (24). Egg clustering within and among kin may increase the local population density of offspring and thus the potential not only for competition but also cooperation at early life-history stages. Benefits from cooperation could include reduced energy expenditure for ovipositing females by minimising search times, reduced egg predation risk (the dilution effect’ hypothesis (25)), increased protection from abiotic challenges, or increased larval communal feeding.

However, cooperative interactions among non-kin can be exploited by cheater individuals. Potential opportunities for cheating can occur in egg-laying species in which mothers provide their eggs with diffusible defensive compounds. One example of defensive compounds is found in the medfly (*Ceratitis capitata*) (26); females provision the surface of their eggs with antimicrobial peptides, which spread into the surrounding environment, creating a zone of microbial growth inhibition. Other examples include in the Green lacewing moth (*Ceraeochrysa cubana*), in which eggs are endowed with an anti-predator alkaloid (27), and in fruit flies (*Drosophila melanogaster*), in which eggs are coated with anticannibalism compounds (28). In each case, these defensive substances represent potential public goods (29, 30) as all nearby eggs could benefit, including those not provisioned with defensive compounds. This raises the intriguing possibility that in egg clusters of mixed maternity, there could be cooperator eggs coated with defensive compounds, and non-defended cheater eggs that benefit from their placement within the defensive diffusion radius. In this scenario, cheaters might benefit by avoiding the energetic expense of provisioning eggs with defensive compounds. Egg clustering might also provide benefits as well as cheating opportunities through communal feeding. For example, if eggs are laid on hard substrates there could be benefits for all via the collective processing of food, while simultaneously allowing the possibility for cheaters to benefit by utilising resources liberated by the processing of food by their cluster mates. Such co-operators and cheats could stably coexist within a population if the benefits of public goods have a non-linear relationship with fitness (30, 31).

The potential fitness benefits of joining an existing egg cluster within a food patch could also depend on key factors such as number, quality, fertility, and age of eggs already present. Based on existing evidence that females can vary the number of eggs they lay according to social density (9, 10, 32) we predict, and test here in the *D. melanogaster* fruit fly model system, the idea that females should use number of eggs and/or number of adult conspecifics as cues to direct their egg-clustering patterns.

### *Drosophila melanogaster* as a model for understanding plasticity in oviposition

In wild populations, *D. melanogaster* larvae and adults periodically occur at high densities around fallen, fermenting fruit. For females, fruit is both a source of nutrition and potential oviposition sites. In *Drosophila* and other species in which fruit resources are patchy, ephemeral, and vary in nutrient availability (density, ripeness, and state of decomposition) access to oviposition sites is likely to be shaped by the immediate social environment (number of conspecific or heterospecific males and females utilising the same resource) (33). To oviposit in a way that maximises fitness, females should assess and respond appropriately to the nutritional quality, degree of interspecific competition, risk of pathogens and parasites present at each oviposition site, as well as the ongoing search costs, and potential benefits and costs for developing offspring (34).

Consistent with this reasoning, previous research shows that female *Drosophila* can show attraction to conspecifics that are utilising specific oviposition resources (35, 36), which they can detect directly or via pheromones, markings, or the presence of eggs or larvae (37–41). Several studies show that gravid females can aggregate to lay their eggs on substrate patches used by others according to a variety of physical and social environmental conditions (9, 42–44). This attraction could arise because it allows females to copy the site-selection choices of others (37, 38, 40) to enable cooperative feeding among larvae (45), or to gain other potential public goods benefits. Existing evidence also suggests that female *Drosophila* can adjust the number of eggs they lay according to their previous or current social environments, laying fewer eggs after exposure to conspecifics before mating (9, 46) but laying eggs more quickly if they are in larger social groups after mating (32).

*Drosophila* eggs can often be found in extremely close proximity, in large clusters (Fig. S1). Hence, after deciding to lay on substrates with existing conspecific eggs, females must also decide whether to lay eggs that join existing clusters. Females generally lay one egg at a time (47), explore oviposition sites between laying, and can retain eggs until an optimal laying site is found (48). Therefore, clustering of eggs suggests that females make repeated egg-laying decisions and that females lay their eggs in non-random distributions. Whether the temporal or spatial distribution of egg laying is significantly non-random, how laying location choices vary *within* a patch, and whether the factors that drive female egg laying patterns and choices vary *within* patches are not known. The pattern of egg laying within patches can be examined by assessing egg clustering patterns, which we define as two or more eggs in direct contact (Fig. 1). Egg clustering is expected to be important, as it offers females the opportunity to optimise fitness by placing their eggs in manner that maximises benefits of cooperation and the potential for public goods benefits (e.g., protection, defence, or cooperative tunnelling) against potential costs of competition (e.g., over-exploitation of food). Clustering by our definition, also potentially enables further benefits only transferred via eggs being in direct contact.

**Figure 1.**
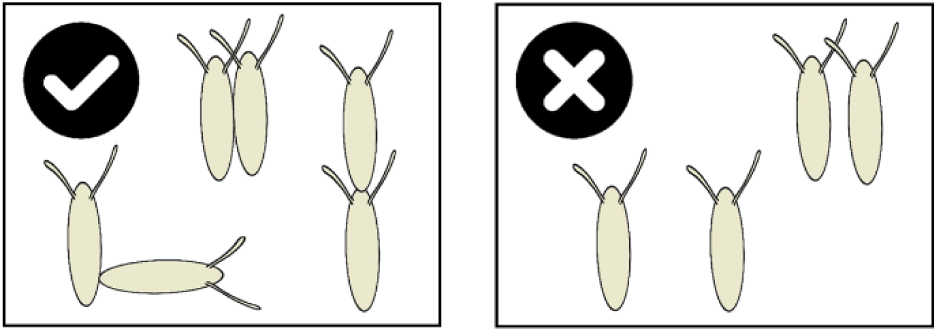
Egg clustering is defined as eggs in direct contact. Eggs where the main bodies were in direct contact (left) were classified as clustered, and those not in contact (or where respiratory appendages only overlap) were classified as singly laid (right).

We tested these ideas with three hypotheses:

1. that female *D. melanogaster* lay their eggs non-randomly within a food patch
2. that changes to adult social density, and the presence of existing egg clusters, alter egg-clustering patterns
3. that females would lay eggs in clusters of mixed maternity at high frequency

Though our initial observations suggests that egg laying patterns *within* patches are non-random, this has not yet been formally confirmed. Therefore, we first constructed a mathematical model, parameterised by experimental data, to test the hypothesis that females exhibit non-random egg placement (H1). We then conducted additional experiments to determine key drivers of egg laying patterns observed. We examined whether egg-clustering patterns responded plastically to adult female group size, or to the density of eggs already present in the local environment (H2). Potential advantages of site copying and egg clustering (9, 17, 18, 25–28, 32, 38, 45, 49, 50) predicted that females should increase egg clustering in response to higher densities of adult females and already-laid eggs in the environment. Females could potentially maximise public goods-related benefits under increased clustering by laying eggs alongside those of others. To establish whether conditions for this scenario exist, we examined whether females so indeed cluster eggs within patches with those of conspecifics by using dyed eggs to distinguish eggs laid by different females (H3).

## Results

We first constructed a model, parameterised with data, to test the hypothesis that females exhibit non-random egg placement (H1). Having confirmed that egg laying patterns were significantly non-random, we then conducted experiments to test the effect of social group size, and presence of existing egg clusters, on the rate and likelihood of egg laying, egg clustering patterns and fitness (egg-adult viability as a function of egg clustering (H2)). The final experiment was then conducted using females that produced dyed eggs, to measure the frequency of egg clusters with mixed maternity (H3).

### 1. Egg laying patterns were significantly non-random, and clustering increased with increasing group size (H1, H2)

Across the four social treatments, clustering preferences (values of К that, based on the Kolmogorov-Smirnov test, yielded no significant difference between the simulated and observed distributions) were between 0.3 and 0.5 (Table S2).

For all social treatments with two or more females, patterns of egg clustering in the empirical data were significantly more clustered than those predicted by the null model (H1; Wilcoxon Signed Rank test comparisons with simulations using К = 0; paired: z_(6)_ = -2.29, p = 0.0220, r = 0.866, N = 7 vials; groups of four: z_(25)_ = -4.47, p = 7.95 x 10^-6^, r = 0.876, N = 26 vials; groups of eight: z_(27)_ = 4.63, p = 3.63 x 10^-6^, r = 0.875, N = 28 vials). Therefore, egg-clustering patterns of all grouped females followed a non-random distribution (Table S2, Fig. 2) but those of solitary females did not (z_(2)_ = -1.36, p = 0.174, r = 0.786, N = 3 vials), suggesting that their laying patterns were random (though we note that parameters were more difficult to estimate for the solitary social treatment as clustering was observed only infrequently under those conditions). Clustering preferences increased with increasing female group size: solitary and paired females had a significantly lower clustering preference in comparison to females housed in groups of 4 or 8 (Table S2, Fig. 2). The findings support H1, that eggs were laid in non-random patterns that became more non-random as social density increased. Therefore, egg clustering patterns were plastic and responsive to the female’s social environment.

**Figure 2.**
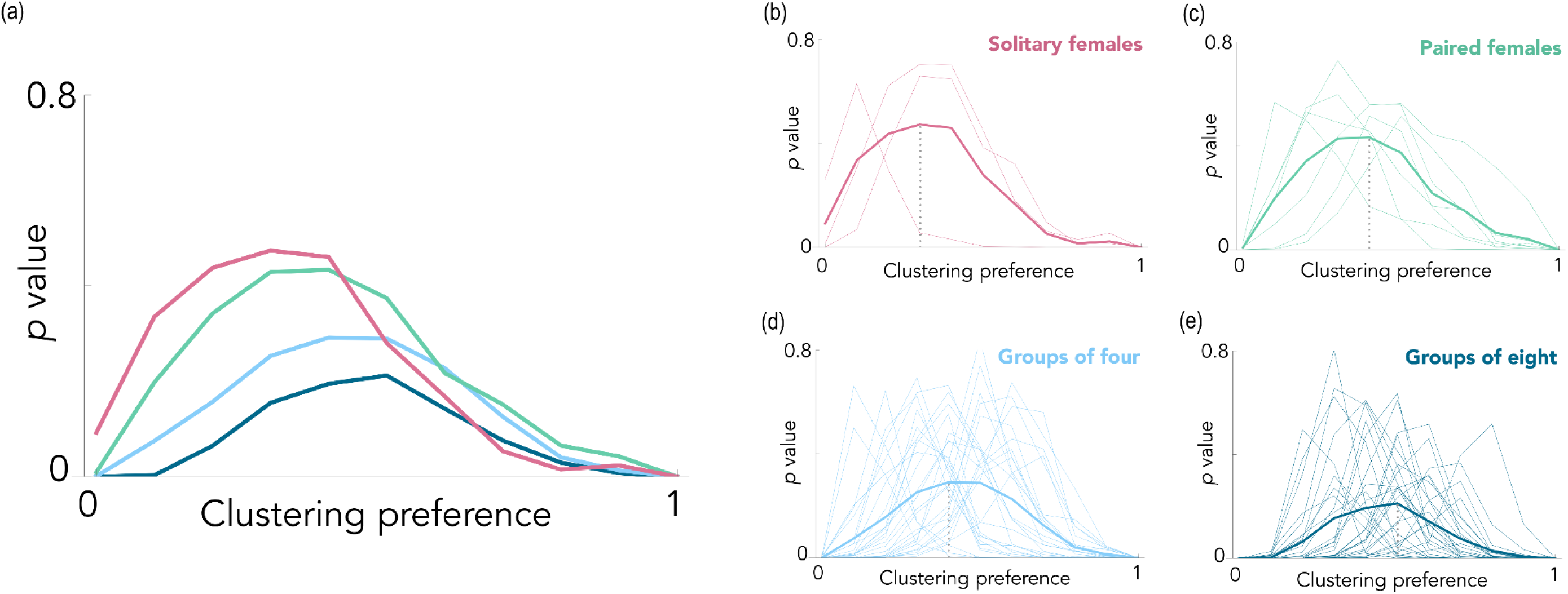
Egg clustering is non-random and increases with increasing adult social density. (a) Comparison of the clustering proportion model outputs and empirical egg laying patterns observed in *D. melanogaster,* for four social group sizes (1, 2, 4, and 8 adult females per group). The average Kolmogorov-Smirnov P values at К (clustering preference) values ranging from 0 to 1 in 0.1 increments – where the null distribution is К = 0 (no eggs clustered), and К = 1 (all eggs laid in one large cluster). The best-fit values of К are those that produced the least significant (highest P value) difference between simulated and observed data. (b-e) The Kolmogorov-Smirnov P values for individual replicates in the four social group sizes (1, 2, 4, and 8 adult females), with the values for each vial shown in the pale, thin lines and the average value indicated with a bold line. The vertical grey dotted lines show clustering preference values.

### 2. Egg-clustering patterns responded both to the density of adult females and presence of already laid eggs in the environment (H2)

#### (i) Females in larger social groups clustered their eggs more and, as expected, laid more eggs and laid eggs at a faster rate

Using the empirical data shown in Fig. 2, we found that females in larger social groups were significantly more likely to lay eggs (X^2^ = 20.2, d.f. = 116, p = 1.51 x 10^-4^). Eggs were present in 100% of vials housing eight females, a significantly higher proportion than for vials containing a solitary female (66.7%, X^2^ = 15.9, d.f. = 58, p = 6.76 x 10^-5^) or those with two females (83.3%, X^2^ = 7.39, d.f. = 58, p = 0.00657). There were also significantly more vials containing eggs from the four female treatment (96.7%) in comparison to the solitary female treatment (X^2^ = 10.2, d.f. = 58, p = 0.00140).

Females in larger social groups also laid more eggs per female than did the females from the other treatments (solitary females: 7 ± 10 eggs per female (N = 30 vials); pairs: 9 ± 9 eggs per female (N = 30 vials); groups of four: 18 ± 8 eggs per female (N = 30 vials); groups of eight: 19 ± 5 eggs per female (N = 30 vials); F_3, 116_ = 265, p = 2.93 x 10^-8^; Fig. 3a). Consistent with this, post-hoc tests showed that females in solitary and paired social densities laid fewer eggs than females in groups of four and eight (all p << 0.001). To control for the increased sampling effort in treatments with larger groups of females, the analysis was repeated with a randomised subset of data to make number of flies (rather than the number of vials) in each treatment similar. The results of these analyses were comparable with the main analysis (larger social groups showed significantly higher likelihood of egg laying: X^2^ = 50.0, d.f. = 54, p = 0.0271 and significantly higher numbers of eggs laid: F_3, 54_ = 137, p = 0.00427).

**Figure 3.**
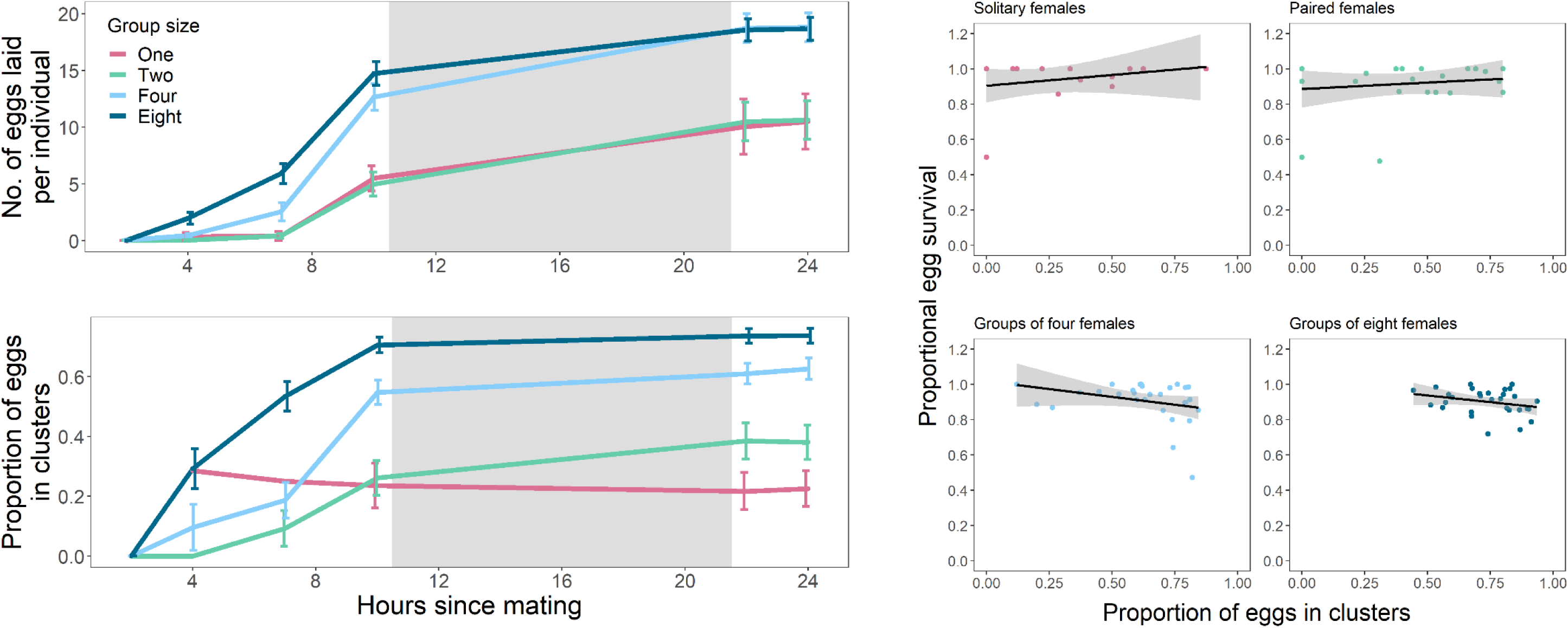
Females in larger groups laid more eggs, more quickly, with a higher proportion of clusters. (a) Females housed in groups of four and eight were quicker to lay, and laid more eggs per female, than those kept in solitude or in pairs (all N = 30 vials). Eggs laid per individual were calculated by dividing total number of observed eggs by the number of females in the vial. Social treatment means and standard errors are shown for the four housing densities at each observation time point. The grey box indicates period of dark (21:00 – 09:00 GMT). (b) Females housed in groups of four or eight laid a higher proportion of their eggs in clusters compared to those kept in solitude or in pairs (all N = 30 vials). Proportion of eggs clustered was calculated by summing all cluster sizes and dividing by the total number of eggs counted. (c) There was no effect of the proportion of egg clustering on egg-adult viability (linear regressions with 95% confidence intervals; solitary: N = 20 vials; paired: N = 24 vials; groups of 4: N = 29 vials; groups of 8: N = 30 vials).

An analysis of the data at the final timepoint (24h post-mating) supported the clustering analysis described in the previous section. Females housed in larger groups laid a higher proportion of their eggs in clusters (solitary females: 22.6% ± 26.9% (N = 20 vials); pairs: 38.1% ± 28.9% (N = 25 vials); groups of four: 62.7% ± 19.2% (N = 29 vials); groups of eight: 73.7% ± 13.3% (N = 30 vials); total number of eggs laid included in model: F_3, 99_ = 28.4, p = 2.53 x 10^-13^; all pairwise comparisons p < 0.05; Fig. 3b). We included the total number of eggs laid in the model as we found that there was a positive relationship between the number of eggs and the proportion of eggs clustered (F_1, 99_ = 6.66, p = 0.0113), although this was not the key driver of social density effects on clustering proportions. In all grouped female treatments, the proportion of clustering also increased over time (individual vial included in model as a random effect: pairs: F_1, 58_ = 19.1, p = 5.26 x 10^-5^; groups of four: F_1, 95_ = 75.8, p = 9.99 x 10^-14^; groups of eight: F_1, 111_ = 74.7, p = 4.45 x 10^-14^). There was no significant effect of time on the proportion of clusters in eggs laid by solitary females (F_1, 40_ = 1.15, p = 0.289).

Consistent with this, the number of clusters and size of the largest cluster increased with both increasing social density (number: F_3, 376_ = 463, p = 2.20 x 10^16^; size: F_3, 299_ = 58.3, p = 2.20 x 10^-16^; Figs. S3-4), and time (number: F_1, 376_ = 631, p = 2.20 x 10^-16^; size: F_1, 299_ = 67.1, p = 7.62 x 10^-15^). These results were robust to differences in sampling effort as shown by an additional analysis using the same number of individuals (rather than vials) at the final timepoint. The effects remained: females in larger groups laid a higher proportion of their eggs in clusters (F_3, 41_ = 6.40, p = 0.00117) and the number and size of clusters still increased with increasing social density (number: F_3, 134_ = 199, p = 2.20 x 10^16^; size: F_3, 102_ = 25.1, p = 2.96 x 10^-12^).

Interestingly, there was no evidence for a fitness benefit for clustered eggs in terms of increased egg-adult survival (Fig 2c). There was no effect of social density (1, 2, 4, or 8 females) on the proportion of eggs that survived to adulthood (F_3, 99_ = 0.292, p = 0.831; Fig. S5) and there was no correlation between the proportion of eggs laid in clusters and egg-adult offspring viability (t = -0.485, d.f. = 101, p = 0.629; Fig. 3c).

Overall, the results supported the hypothesis that the egg-clustering behaviour of females within a food patch responds plastically to the social environment, with females laying eggs more quickly, laying more eggs and laying more eggs in clusters, as the social group size increases. However, there was no indication that eggs laid in clusters had higher egg-adult viability.

#### (ii) Females preferred to lay eggs within existing clusters rather than with singly laid eggs

Female egg-clustering behaviour also showed plasticity in response to eggs already present in the environment. Females were more likely to add at least one of their eggs to existing clusters of 4, 7, or 10 eggs, than they were to lay with a singly placed existing egg (X^2^ = 12.5, d.f. = 156, p = 4.18 x 10^-4^; Fig. 4a). The total proportion of eggs laid that joined existing eggs was also higher when existing eggs were clustered (F_3, 154_ = 4.75, p = 0.00340; Fig. 4b). Proportions were calculated from the number of eggs females were able to lay within the 30 min observation window (between 1 – 19 eggs per vial). Post-hoc tests showed that egg cluster sizes of 4 and 10 were joined significantly more often than were single eggs (four: F_1, 71_ = 13.0, p = 5.85 x 10^-4^; ten: F_1, 78_ = 8.76, p = 0.00407), though this trend was marginally non-significant for a cluster size of 7 (F_1, 75_ = 3.82, p = 0.0542).

**Figure 4.**
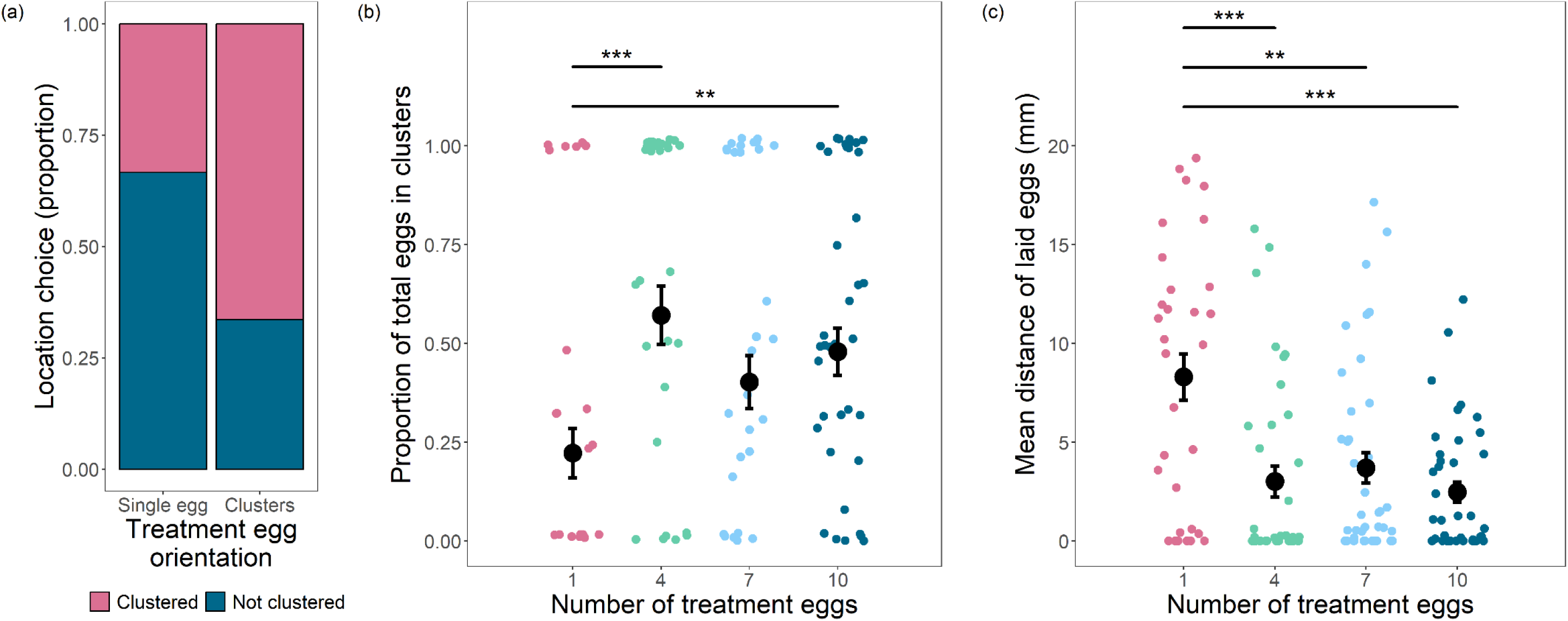
Females lay more eggs within existing clusters and lay eggs closer to existing egg clusters. (a) A higher proportion of females clustered at least one of their eggs with existing eggs, when those existing eggs were in clusters compared to singly laid eggs (singly laid eggs: N = 36; clustered eggs: N = 122. (b). Females laid a higher proportion of their eggs in clusters with existing eggs, when those existing eggs were already in a cluster (of any size) compared to a singly laid egg (1 egg: N = 36; 4 eggs: N = 37; 7 eggs: N = 41; 10 eggs: N = 44). Means and standard errors are shown for the four treatment egg cluster sizes. Significant differences between treatment egg cluster sizes are represented by an overarching bar; * indicates a significant difference between paired treatments (* P < 0.05, ** P < 0.01, *** P < 0.001). (c). Mean distance of laid eggs from existing treatment eggs was greater when the number of existing eggs was one, compared to egg clusters of any size (1 egg: N = 32; 4 eggs: N = 37; 7 eggs: N = 40; 10 eggs: N = 40).

In addition to scoring whether a laid egg was physically in contact with an existing egg (i.e. clustered), we also measured the distance between each newly-laid egg and existing treatment eggs already present on the substrate. We did this to test for potential benefits of close-proximity in addition to direct contact. Females laid their eggs closer to existing clusters (H3; F_3, 145_ = 9.86, p = 5.86 x 10^-6^; Fig. 4c) with all three cluster sizes having shorter inter-egg distances than eggs from females exposed to single eggs (four: F_1, 67_ = 14.8, p = 2.68 x 10^-4^; seven: F_1, 70_ = 11.4, p = 0.00122; ten: F_1, 70_ = 23.8, p = 6.57 x 10^-6^). Hence, laid eggs that did not strictly join a cluster were still laid closer to existing clusters than to single eggs. The results show that females also respond plastically to the clustering patterns of eggs already present in the environment and, consistent with the results above, lay their eggs preferentially within existing egg clusters (H3).

### 3. Egg clusters were typically of mixed-maternity (H3)

In the final experiment we used a lipophilic dye to enable us to distinguish between eggs laid by different females (51). We set up groups of 4 females comprising 1 standard-fed focal wild type and 3 Sudan Black B dye-fed non-focals. We found that of the focal eggs that had been laid in clusters, a mean of 79.5% of them were in mixed-maternity clusters (Fig. 5). This showed that females do not preferentially choose to isolate their eggs by laying them solely in single-maternity clusters, but instead lay eggs in clusters of mixed-maternity at high frequency.

**Figure 5.**
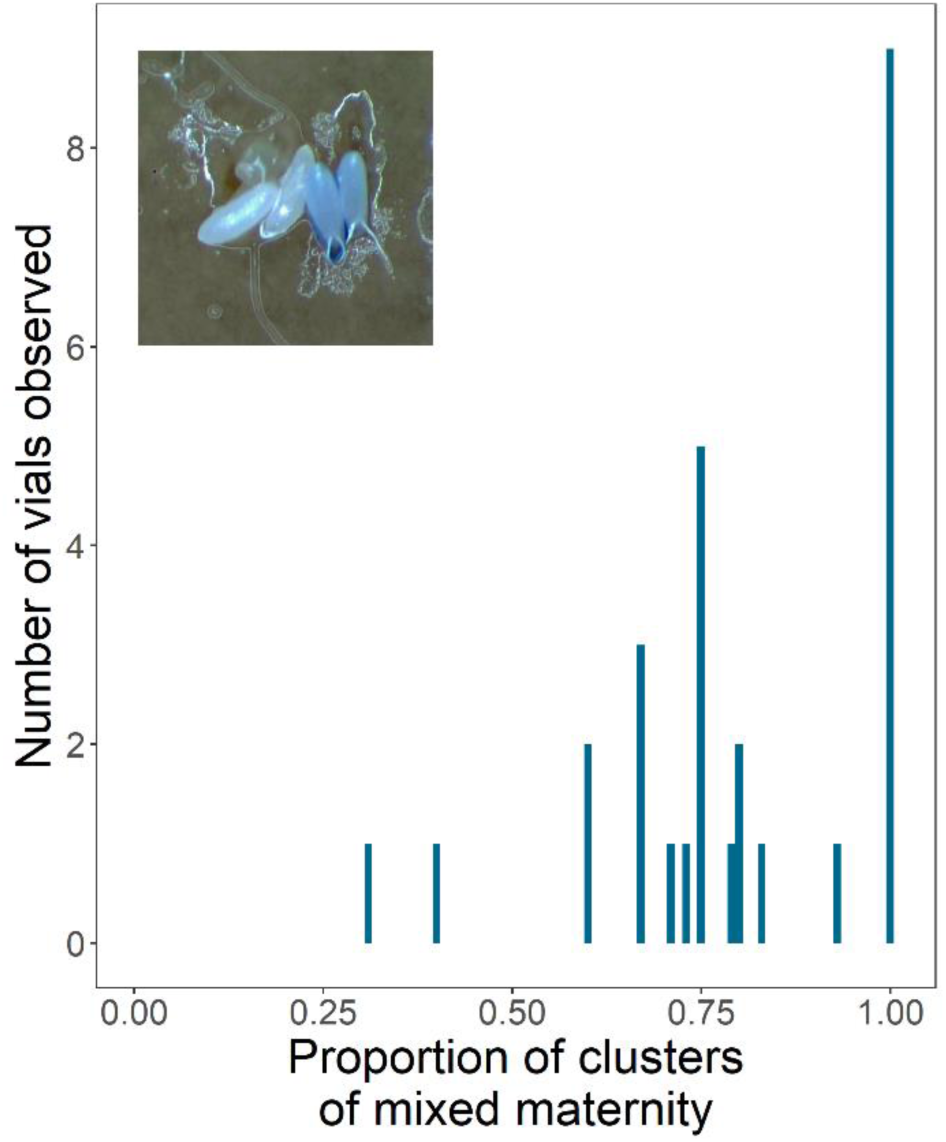
High frequency of mixed maternity egg clusters. Females showed no preference for laying their eggs in single-maternity clusters, as 79.5% of egg clusters observed were of mixed maternity (N = 28 vials). Shown is a frequency distribution of the proportion of egg clusters counted that had eggs laid by two or more gravid females. The inset picture shows a cluster of four eggs: two undyed and two dyed with Sudan Black B dye.

## Discussion

The main new findings of this study were that female egg-clustering behaviour within patches responded plastically to the social environment and to the presence of existing eggs within the oviposition environment. All three hypotheses were supported. 1) Egg-clustering patterns were significantly non-random, for females in groups, but not for eggs laid by socially isolated females. 2) As the social group size of females increased, females laid eggs more quickly, laid more eggs, and laid more of them in clusters. However, there was no evidence of a fitness benefit to clustered eggs in increased egg-adult viability. 3) Clusters of eggs were usually of mixed maternity and females preferred to add their eggs to existing clusters. Overall, females showed striking plasticity in their egg-clustering decisions, with a strong bias towards laying eggs in mixed-maternity clusters. However, the fitness benefits of such behaviour remain elusive. These results are explored in more detail, below.

### Egg-clustering decisions are plastic and eggs are laid in non-random patterns

For all females except those held in social isolation egg clustering occurred more than the chance outcomes predicted by the model. This provides evidence that females make active egg location decisions, and that these responses are plastic and change in response to varying social environments. Egg clustering shows that females make repeated egg-laying decisions, as they generally explore potential laying sites between laying each individual egg, and can retain their eggs until they find an optimal site (47, 48). Increased clustering has the potential to increase local population density within patches, which has potential cumulative effects, as even relatively small-scale variation in density leading to increased competition can have consequences for fitness (52).

### Females cluster more of their eggs in larger groups, and more commonly form mixed-maternity clusters

The proportion of eggs that were laid in clusters increased as social density increased. Although there was a higher chance of clustering when there are more eggs present in the environment, this increase in clustering with increasing social density did not depend upon egg number (in agreement with the null model comparison). It was not possible to determine the maternity of the clusters in grouped treatments in the initial experiment, but our later experiment showed that the clusters are usually of mixed maternity. Females exposed to other (non-kin) females preferred to add their eggs to mixed-maternity clusters, and overall socially isolated females cluster their eggs much less, which suggests that there are potential benefits of laying in mixed-maternity clusters over and above laying in clusters *per se*. The drivers of this are not yet known, however females could be balancing a bet-hedging strategy of laying in multiple sites to reduce risks of sibling competition (if food is limited) or to minimise predation/parasitism risk by reducing transmission and egg visibility (53), whilst adding to existing clusters to reap potential public goods benefits of acquiring more diverse microbiomes (54, 55) or diffusible defensive compounds such as anticannibalism pheromones (28) or antimicrobials (as seen in medfly (26)), but see (51). Mixed-maternity egg clustering could select for cheating among females under a public goods system (29, 30), a possibility that would be interesting to test further.

### Females preferred to lay their eggs in existing egg clusters

Females preferred to add their eggs, or lay them close to, existing clusters of any size. Interestingly, the strength of this preference did not increase with increasing cluster size. This could be because (i) there is no benefit to increasing cluster size (i.e., benefits to joining clusters are not additive), (ii) the clusters here did not vary sufficiently in size, or (iii) there is some trade-off to the benefits of clusters of these sizes. For example, eggs laid later may be at a higher risk of cannibalisation (28, 56) and the risks and costs might increase with cluster size. Costs of laying in mixed-maternity clusters are also possible if being surrounded by kin provides benefits via communal feeding of larvae (13). Exactly how individuals might identify maternity of egg clusters within patches, whether by detecting conspecific pheromones (41, 57, 58)) or absolute cluster size remain to be investigated. The strong aversion to laying with isolated eggs raises the question of how clusters are initiated in the first place. Female choice to copy and lay with other eggs could require a threshold of a repeated decision – i.e., at least two eggs at the same site, because this is a more robust indicator of site quality than only one. Females may avoid laying with existing single eggs if those have a higher probability of infertility and thus offer potentially fewer benefits. However, we found no relationship between cluster size and egg-to-adult viability, which argues against this idea. In addition, whether females can detect infertile eggs is not known, although those laid by virgins have a distinct pheromonal profile (41)). One scenario is that clusters form if the first two eggs are laid by the same mother. Arguing against this idea is that clusters of two eggs laid by different mothers were observed in our mixed-maternity investigation. This suggests that further studies on clustering initiation are required.

### Potential fitness benefits of egg clustering remain elusive

Interestingly, we detected no increase in egg-adult viability with increasing egg clustering. It is not clear why we did not observe fitness benefits, given potential costs of both maintaining plasticity (59, 60) and of increasing local competition (13, 28, 34, 52–56). It is possible that as mothers were able to choose a clustering strategy, all individuals chose the optimal strategy. To test this, future work could force mothers into making an incorrect choice via a mismatch of environments. It is also possible that the *ad libitum* food provided obscured any costs of choosing the ‘wrong’ egg laying strategy. It was necessary to remove females from the egg laying environments before they had oviposited all fertile eggs, as eggs needed to be counted prior to larvae hatching which takes place after 22 – 24 hours at 25°C (61, 62). Thus, it is also possible that differences in egg-adult viability could have been detected if females were able to continue laying if the benefit only occurs in clusters of mixed-age eggs.

### Egg number and laying rate also increased as adult female density increased

As expected, based on existing research (32), females in larger social groups laid more eggs and laid them more quickly. This is consistent with raffle theory, in which the production of more eggs when competition is high ensures that mothers have a greater chance of at least some of their offspring surviving to adulthood (63). With a greater number of conspecifics present in the environment, there should also be a higher chance of copying behaviours arising, because a greater number of potential ‘initiators’ are present (37, 38, 40). Increased copying could increase laying rates, benefitting mothers as it may reduce sampling times needed to locate optimal oviposition sites. Consistent with this, we observed that latency to lay the first egg was also shorter at higher adult densities. In situations in which larval density is high and resources are limiting, later-laid eggs could have reduced survival due to resource degradation and cannibalisation (13, 32, 56). Thus, in high density environments, mothers should ensure they begin laying quickly to avoid increased offspring lethality.

Previous research (32) showed that the daylight period can inhibit egg laying in *Drosophila*, an effect that can be over-ridden by being in a group (five females). Intriguingly, our results showed remarkably similar laying rates for isolated and paired females (whereas groups of four or eight females matched laying times reported in (32)), suggesting that exposure to only one other gravid female does not counteract the previously described light-induced inhibition of egg production. This is surprising given that in male *Drosophila*, exposure to one other rival is enough to facilitate reproductive behavioural responses (64–66). This may suggest that there are sex differences in responses to local population density.

## Conclusions

Overall, we demonstrated that *D. melanogaster* mothers express plasticity in egg-clustering decisions, laying eggs in a significantly non-random manner and responding to differences in social density from multiple life stages. Exposure to increased adult social density resulted in females laying more eggs, at a faster rate, and, a new result from this study, laying a higher proportion of those eggs in clusters. Exposure to existing egg clusters also led to a higher frequency of clustering decisions, regardless of existing cluster size. We hypothesise that eggs may be clustered to gain public goods benefits. However, the overall ecological importance, and the costs and benefits of this plasticity remain unknown. These results demonstrate that egg laying and clustering decisions are highly sophisticated and add to the growing evidence that even non-social organisms such as *D. melanogaster* have unexpectedly rich social lives (67, 68).

## Methods

### Fly husbandry and rearing of experimental flies

Fly rearing and all experiments were conducted at 25°C on a 12h light:dark cycle (09:00-21:00 GMT). Wild type flies originating from a large Dahomey laboratory population maintained in cages with overlapping generations were used throughout. We collected eggs from cages using Petri dishes (*Sarstedt #82.1473.001*) filled with grape juice agar-based medium (50g agar (*Fisher Scientific #10048991*), 600ml red grape juice (*Young’s Brew red wine enhancer*), 42ml Nipagin solution (methylparaben, 10% w/v solution, dissolved in 95% Ethanol) per 1.1l H_2_O). Once eggs had hatched, we transferred first instar larvae to a 40ml plastic vial (*Sarstedt #58.490*) containing 7ml of a standard sugar-yeast-agar (SYA) medium (100g brewer’s yeast (*Buy Wholefoods Online*), 50g sugar (*Tate & Lyle*), 15g agar, 30ml Nipagin solution, and 3ml propionic acid (*Fisher Scientific #10193190*) per litre of medium), at a standard density of 100 larvae per vial (i.e., ‘standard density vials’). Experimental flies were then collected under light ice anaesthesia, within six hours of eclosion to ensure virginity.

#### 1. Development of mathematical model of egg-clustering to test for non-random egg deposition (H1)

To ascertain whether the proportion of observed egg clustering differed from a null expectation, we produced a model to calculate random distributions of eggs in a standard vial. We defined a ‘cluster’ as any group of two or more eggs in which the main bodies of the eggs were in direct contact (Fig. 1). In the model, given that the observed egg to dish area is 1/5770, we assume that: (i) there are 5770 positions available for egg laying, and (ii) there is a total number (parameter Ɛ) of eggs per vial. Assuming that all eggs are laid sequentially and at random, we calculated the expected size of clusters (defined as the number of eggs per cluster) for any number of total eggs laid (Ɛ).

The preference for clustering eggs next to an existing egg (hereafter ‘clustering preference’) is defined by the probability of a clustering decision. К is the probability that a fly will lay an egg next to an already existing egg. After the first egg is placed at random, where Ɛ > 1, the next egg is either laid in a random position (probability of probability of 1 - К) or next to an existing egg (probability of К); in the latter case, the position of the egg is chosen at random amongst only positions with existing eggs.

The expected number of eggs per cluster can be predicted based on К and Ɛ. When К = 0 (no preference for egg clustering), very few clusters with 2 eggs occur and virtually no clusters with more than 2 eggs (2.5% with Ɛ = 233 (the maximum number of eggs observed in our experiment); 1% with Ɛ = 100; no clusters for Ɛ below 70). Therefore, if Ɛ << 5770 (as is the case in our experiment), randomly laid eggs form almost exclusively singly laid eggs. When К = 1 (absolute preference for egg clustering), all eggs are clustered in one large cluster in a single position.

At intermediate values of К, some variation in cluster size emerges. We simulated the expected number and size of egg clusters for different values of К from 0 to 1 in intervals of 0.1 (using the average of 10 different simulations per К value). We then compared these expected distributions to the empirically observed distributions of eggs laid by females from the differing social density treatments (groups of 1, 2, 4, or 8) described below, to estimate the most likely preference К leading to that distribution. For each vial, we performed a Kolmogorov-Smirnov test (implemented in Mathematica 13. 1 (69)) between the expected and observed distribution, for all К values. We considered the К value yielding the highest (least significant) P values to be the most likely К value for that vial.

#### 2. Testing for plasticity in egg-clustering behaviour in response to varying egg and adult density (H2)

##### (i) Effects of adult social density on egg-clustering decisions and fitness

We investigated how variation in adult social group size affected egg laying and egg-clustering decisions: specifically, the speed of egg laying, the number of eggs laid, egg clustering patterns, and fitness (egg-adult viability of clustered versus non-clustered eggs). Some data from this experiment were also used to parameterise the model above. Experimental flies were collected and housed in same-sex groups of 10 in standard vials. At four days post-eclosion, females were randomly assigned to one of the following social group treatments: 1, 2, 4, or 8 females and kept in these conditions for 48h (N = 30 vials each treatment; Fig. S1). Males were held for six days until they were used for matings.

At 6 days post-eclosion, males were introduced to all the female social group treatment vials for 2h at a density of 3:2 males:females (or 2:1 for isolated females) to enable choice (Fig. S1). To ensure that this was a set up that would result in all females successfully mating within the 2h period allowed, we separately recorded female mating frequency in an identical experimental set up. In this we observed that 97.3% of females mated, with no significant difference in mating frequency between treatments (X^2^ = 1.79, d.f. = 3, p = 0.617). Thus, we can assume that there is only a very low probability of females remaining unmated by using this procedure.

After the 2h mating period in the main experiment, we transferred all mated females to fresh standard vials to observe egg laying decisions. Every 2 – 3h, we counted the number of eggs laid, number of egg clusters, and size of egg clusters. Observations were made at 14:00, 16:00, 19:00, and 22:00 on day 1 of laying, and 10:00 and 12:00 on day 2 (i.e., 2, 4, 7, 10, 22, and 24h after the end of the mating period, respectively). Differences in counts (number of eggs, number of clusters, and size of clusters) between timepoints were calculated to give overall egg laying latencies, numbers and clustering rates. We calculated the proportion of eggs in clusters at each timepoint by summing the total number of eggs in each cluster and dividing by the overall total number of eggs laid in the vial. All females were removed after 12:00 on day 2 and vials were then kept for 14 days, to count the total number of eclosed adults from each of the social exposure treatments.

##### (ii) Effects of existing egg clusters on female egg-clustering decisions

To assess whether the size of egg clusters already present in an environment alters subsequent female egg-clustering decisions, we presented focal females with a laying environment that already contained varying numbers of eggs and egg clusters. To create the egg clusters, singly laid eggs were collected within 2 hours of laying from SYA medium-filled Petri dishes and transferred into vials to create four different egg cluster sizes: 1, 4, 7, and 10 eggs per cluster. Females have a strong tendency to lay eggs at vial edges (Supplementary Information Text; Table S1; (9, 49, 50)), hence we simulated this preference by placing all egg clusters at the edge of the egg treatment vials.

To obtain the Dahomey females and males for the experiment, we collected virgin flies from standard density vials. Prior to the tests, females were maintained alone, males in groups of four per vial. At seven days post-eclosion, males and females were paired and observed to ensure they had all successfully mated. After mating, groups of 4 females were then transferred to each of the 4 types of egg cluster treatment vials (1 egg: N = 20 vials; 4 eggs: N = 17 vials; 7 eggs: N = 20 vials; 10 eggs: N = 21 vials). After the introduction of the focal females, vials were checked at 30 min intervals from 13:00 until 21:30, at which point if females had not laid eggs they were removed. If eggs were laid by the introduced focal females, we counted the number of eggs laid and categorised them in relation to the existing egg cluster present (clustered or not clustered). All vials were then frozen at -20°C for imaging.

Vials were imaged to record inter egg distances, using a video camera (Sony Handycam HDR-CX405). Using ImageJ’s multiple point selector tool (70) we captured x-y coordinates of each egg and converted these into Euclidean pairwise distances. Selected coordinates were taken from the edge points of eggs that were in closest proximity, resulting in clustered eggs having a distance value of zero.

#### 3. Testing for mixed maternity of egg clusters (H3)

In the final experiment, we determined the extent of egg clusters of single or mixed maternity. In each treatment group there was one wild type (Dahomey) focal and 3 non-focal females from the *scarlet* strain (recessive *scarlet* eye colour mutation backcrossed into the Dahomey wild type > 4 times). These non-focal *scarlet* females were fed Sudan Black B dye that stained laid eggs, allowing the eggs of focal versus non-focal females to be identified. First instar larvae of Dahomey wild type and *scarlet* strains were raised in cultures of 100 larvae per vial as described above. Focal Dahomey females were collected from cultures reared and maintained on standard medium, whereas *scarlet* females were derived from cultures containing Sudan Black B dye (100g brewer’s yeast, 50g sugar, 15g agar, 30ml Nipagin solution, 3ml propionic acid, and 1.4g Sudan Black B powder (*Sigma-Aldrich,* cat#199664) dissolved in 14ml corn oil (*Mazola* 100% pure corn oil) per litre of medium). Focal Dahomey virgin females were collected from the standard medium vials, and non-focal *scarlet* virgin females were collected from Sudan Black B vials and maintained on that same medium until use. Upon collection, all females were housed in groups of 10 and then Dahomey focal females were housed in groups of 4 from 5 days post-eclosion. At 4 days post-eclosion, *scarlet* females were wing clipped for identification, and also housed 4 per vial. Dahomey and *scarlet* males for matings were collected from standard medium vials and housed in groups of 10 until use.

Seven days post-eclosion, males were transferred into the female vials for mating: 8 Dahomey males were added to each group of 4 Dahomey females, and 6 *scarlet* males for each group of 4 *scarlet* females. The flies were given 4 hours to mate. One female from each of the Dahomey vials was then transferred to fresh standard media alongside three *scarlet* dye-fed females at 12:00 GMT (N = 29). We then counted the number of eggs laid at 10.5h post-mating. Eggs were categorised as clustered or not clustered (Fig. 1) and according to whether they were laid by a focal or non-focal female. This allowed us to calculate the proportion of eggs that were clustered, and whether the cluster contained eggs from both focal and non-focal females.

### Statistical analysis

All statistical analyses were conducted using R v 4. 2. 2 (71), and mean values and standard deviations are reported within the text. Graphs were produced using packages ‘ggplot2’, ‘cowplot’ and ‘grid’ (72, 73), with plotted standard errors calculated using ‘Rmisc’ (74).

We analysed the effect of egg and adult social density on the likelihood that females laid eggs and produced adult offspring, by using generalised linear models (GLMs) with binomial errors. We used the package ‘AER’ (75) to test for overdispersion, and where that was found, we used GLMs with a quasi-Poisson distribution to account for this over-dispersion and analyse the effect of adult density on the number of offspring produced. Linear models were used to investigate the effect of adult density on egg-clustering patterns (egg laying and clustering rates), and total eggs laid was included as a fixed effect to account for increased likelihood of clustering when more eggs are present in the environment.

When analysing variables throughout multiple timepoints, we used linear mixed effects models; the timepoint was included as a fixed effect in the model to account for differences in rates and the test vial was included as a random effect – using functions in packages ‘lme4’ (76) and ‘lmerTest’ (77). We tested for Pearson’s product-moment correlations between the proportion of eggs laid in clusters and subsequent egg-adult viability. In all cases, post-hoc tests (models including only pairwise comparisons) were used to identify significant differences in the six pairwise comparisons available.

In the first experiment investigating the effects of social density, there was variation in sampling effort between treatments, due to the differences in female group sizes (and lack of identification of focal females). To account for this, all significant results from the initial analyses were reanalysed with a randomised subset of data so that the number of individual flies (rather than number of vials) per treatment were similar. In this reduced subset, all vials from solitary females were included (N = 30 females), half of those in pairs (N = 30 females), eight of those from groups of 4 (N = 32 females), and four of those from groups of 8 (N = 32 females). These analyses were congruent with those of the initial analyses, suggesting the initial analyses were robust.

## Declarations

### Ethics approval and consent to participate

Not applicable.

### Consent for publication

Not applicable.

### Availability of data and materials

The datasets generated and/or analysed during the current study are attached for review as supplementary material and will be openly available from the Leeds Research Data Repository on publication.

### Author’s contributions

*Conceptualisation*: Emily R. Churchill, Emily K. Fowler, Lucy A. Friend, Marco Archetti, Douglas W. Yu, Andrew F.G. Bourke, Tracey Chapman, and Amanda Bretman. *Data curation*: Emily R. Churchill, Emily K. Fowler, and Lucy A. Friend. *Funding acquisition*: Emily K. Fowler, Marco Archetti, Douglas W. Yu, Andrew F.G. Bourke, Tracey Chapman, and Amanda Bretman. *Investigation*: Emily R. Churchill, Emily K. Fowler, Lucy A. Friend, and Marco Archetti. *Methodology*: Emily R. Churchill, Emily K. Fowler, Lucy A. Friend, and Marco Archetti. *Project administration*: Emily R. Churchill, Emily K. Fowler, Lucy A. Friend, Tracey Chapman, and Amanda Bretman. *Resources*: Tracey Chapman and Amanda Bretman. *Software*: Emily R. Churchill and Marco Archetti. *Visualisation*: Emily R. Churchill and Marco Archetti. *Writing – original draft*: Emily R. Churchill. *Writing – review and editing*: Emily R. Churchill, Emily K. Fowler, Lucy A. Friend, Marco Archetti, Douglas W. Yu, Andrew F.G. Bourke, Tracey Chapman, and Amanda Bretman. All authors gave final approval for publication.

## Acknowledgements

We would like to thank Paul Candon and Kerri Armstrong for technical assistance, and all anonymous reviewers of this manuscript for their thoughtful and constructive feedback, which great enhanced the quality of this manuscript.

## Supplementary Information

### Supplementary text 1. Determination of effect of natural egg placement and egg ‘edge effects’ on subsequent egg laying behaviour

In this study, we tested the responses of females to the presence of existing eggs of various cluster sizes in the environment. This required us to manually place eggs in vials to create the required variation in egg clustering. Previous research shows that females naturally prefer to lay eggs at the edges of substrates (9, 47, 48). Such edge biases could themselves influence the decisions of subsequent females about whether to create egg clusters. To test for this, and thus understand how best to place eggs in the main experiment, we conducted an initial preliminary experiment to test whether proximity of existing eggs to the edge off the vial affected the subsequent latency of females to lay eggs near them.

We tested isolated versus grouped females laying eggs in 3 types of ‘egg treatment’ vials: in which there was (i) no egg present (control), and (ii) 1 egg either placed in the centre or (iii) at the edge of the vial. The non-focal eggs used to create the egg treatment vials were laid by groups of 10 gravid females allowed to lay overnight on Petri dishes filled with 40ml of SYA medium. To standardise eggs, we selected only eggs laid naturally in isolation from these dishes for subsequent transfer to create the egg treatment vials. To enable transfer of eggs without damage to the SYA medium, elected eggs were transferred using a mounted needle, into standard plastic vials that had been cut in half. Then the top half was reattached after egg transfer using Sellotape. Test flies were collected from standard density vials (as in the main experiments) and unmated females were housed in solitude, and males in groups of four. After six days, one male was transferred into the vial with each female, and pairs were watched to ensure matings occurred. We then transferred females in isolation or in groups of 4 into vials containing the 3 egg treatments (no egg, 1 edge egg, or 1 central egg; N = 15 - 42 vials per treatment).

We then checked vials every 30 minutes until 21:30 (one count after lights off in the constant environment room) and removed the female(s) if they had laid eggs. If they did not lay, they were kept overnight, and egg counts resumed at 09:30 for three further hours, until 24 hours after females had initially been introduced into the vials. We calculated egg laying latencies and the number of eggs laid by the focal females. We also noted whether eggs were laid in a cluster (defined in Fig. 1) alongside the existing egg.

The results revealed a strong preference to lay at the edge of the vial, confirming previous reports. There was no effect of existing egg location (X^2^ = 1.32, d.f. = 113, p = 0.517) or group size (X^2^ = 3.80, d.f. = 112, p = 0.0513) on where in the environment females subsequently laid their eggs. There was also no effect of either egg location or group size on whether females chose to cluster their eggs as no females clustered their eggs (Table S1). Females housed in solitude took longer to initiate laying than did those in groups of four (X^2^ = 4.38, d.f. = 126, p = 0.001; Fig. S2). There was no effect of existing egg location on laying latency (X^2^ = 1.30, d.f. = 125, p = 0.139; Fig. S2). Based on these results, we created egg clusters in the main experiments by following the natural patterns by placing eggs at the vial edges.

**Figure S1.**
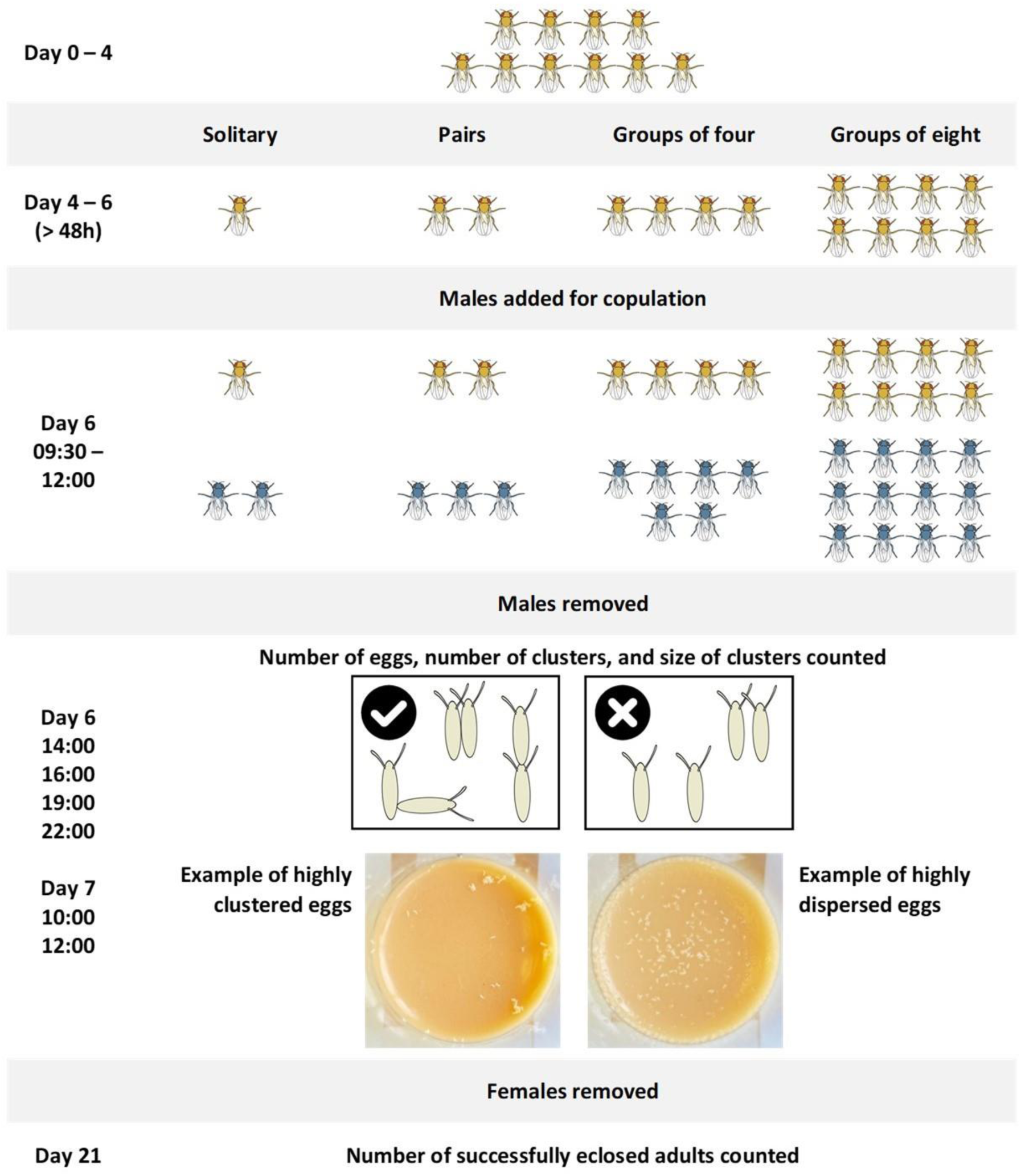
Experimental set up for testing the effects of social densities on egg laying. Summary of the experimental set up for testing the effects of social densities on egg laying. Day 0 – 4: males and females housed in same-sex groups of 10. Day 5 – 6: females transferred to one of four social treatments (whilst males remained in groups of 10). Day 6: males were added to female treatment vials in a ratio of 3:2 (2:1 for isolated females) and given 2h to mate. Day 6 – 7: number of eggs, number of clusters and size of clusters were counted at the six listed timepoints. Shown is the strategy for defining egg clusters (contact between the main body of eggs).

**Figure S2.**
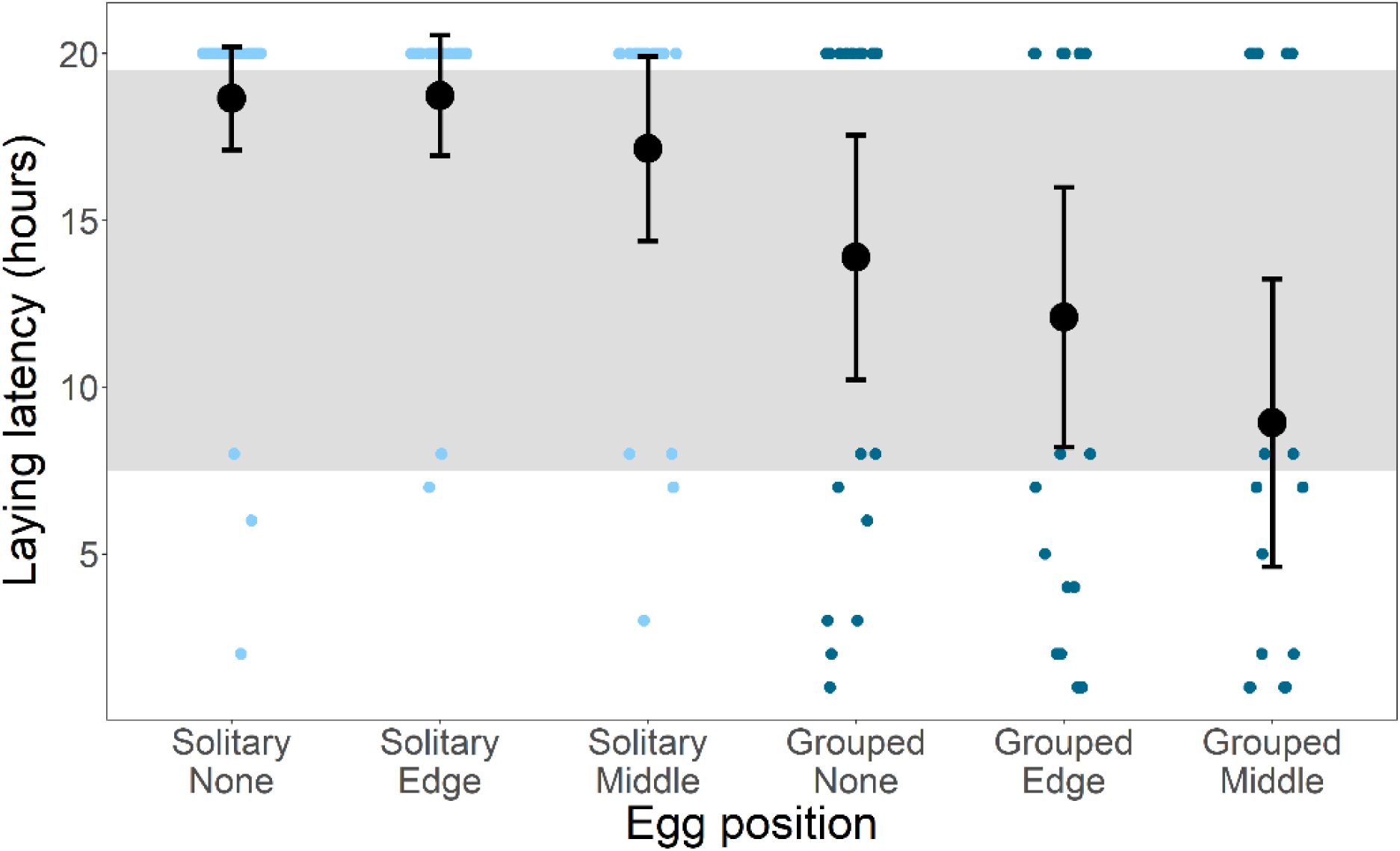
Existing egg location did not impact latency to lay. Females housed in solitude were slower to start laying than those housed in groups of 4, but there was no effect of existing egg location on laying latency. Means and standard errors are shown for the six treatments. Grey box indicates period of dark (21:00 – 09:00 GMT). N = 30 vials for all four social treatments.

**Figure S3.**
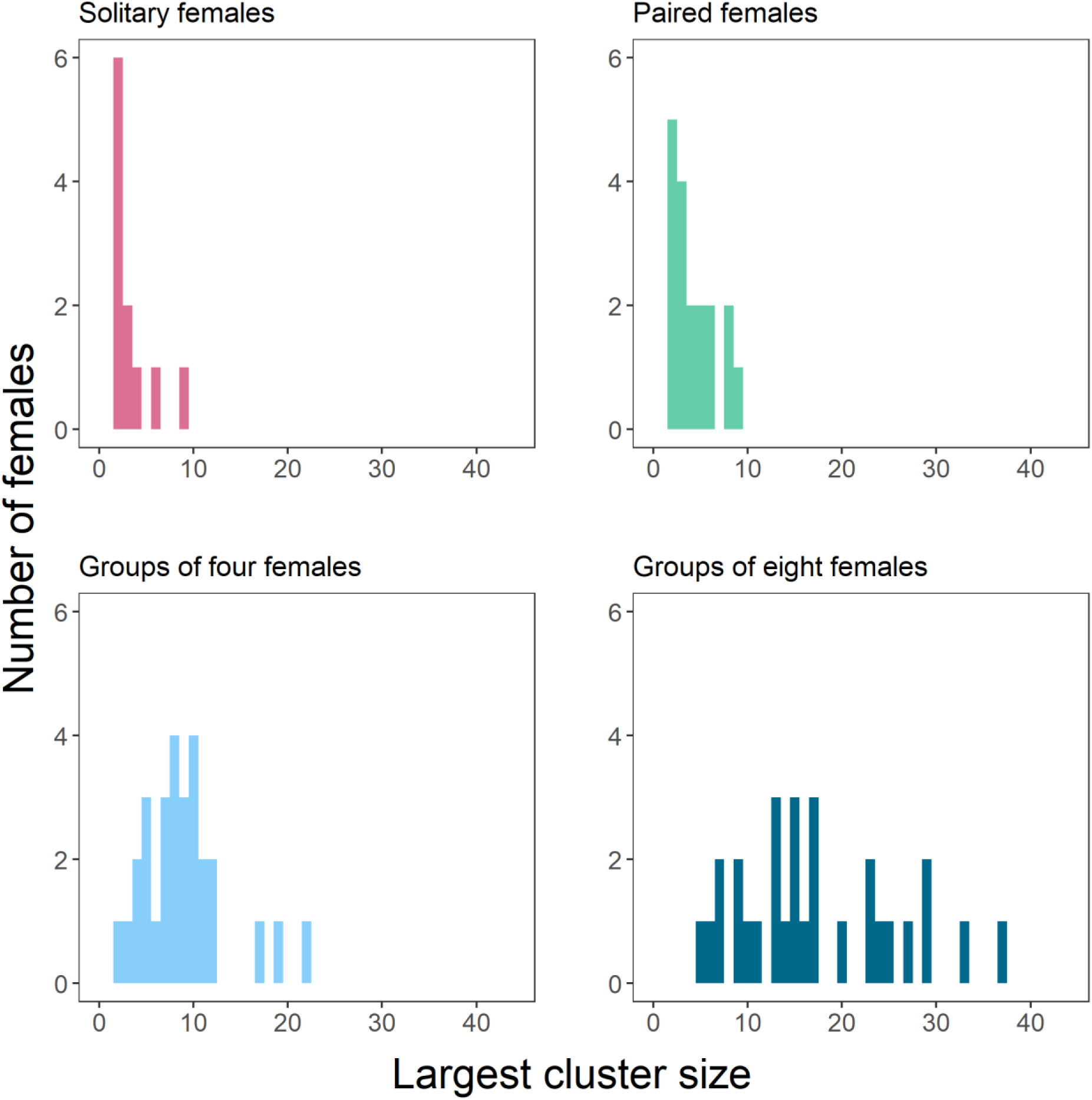
Females in larger social groups laid significantly larger clusters. Maximum cluster size increased with increasing female social density. Shown is a frequency distribution of the largest egg cluster for mated females placed in social groups of 1, 2, 4, and 8 for 24 hours. Clusters were defined as in Fig. 1. Data are shown as the total number of vials with each cluster size observed for each of the four treatments (solitary females: N = 11; pairs: N = 18; groups of 4: N = 29; groups of 8: N = 30 vials).

**Figure S4.**
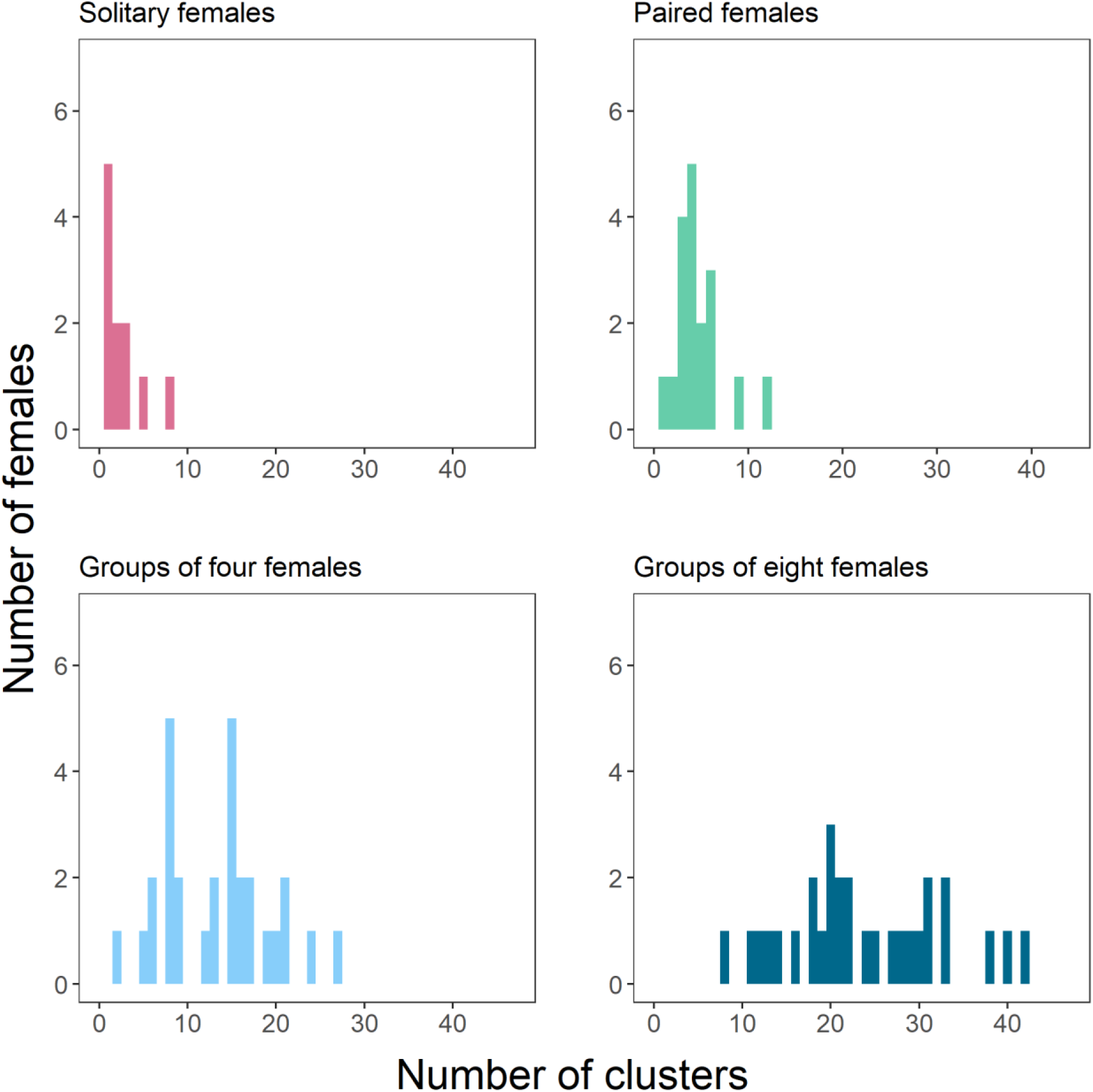
Females in larger social groups laid significantly more clusters. A greater number of clusters were observed with increasing female social density. Shown is a frequency distribution of the egg clustering sizes for mated females placed in social groups of 1, 2, 4, and 8 for 24 hours. Clusters were defined as in Fig. 1. Data are shown as the total number of vials with each cluster size observed for each of the four treatments (solitary females: N = 11; pairs: N = 18; groups of 4: N = 29; groups of 8: N = 30 vials).

**Figure S5.**
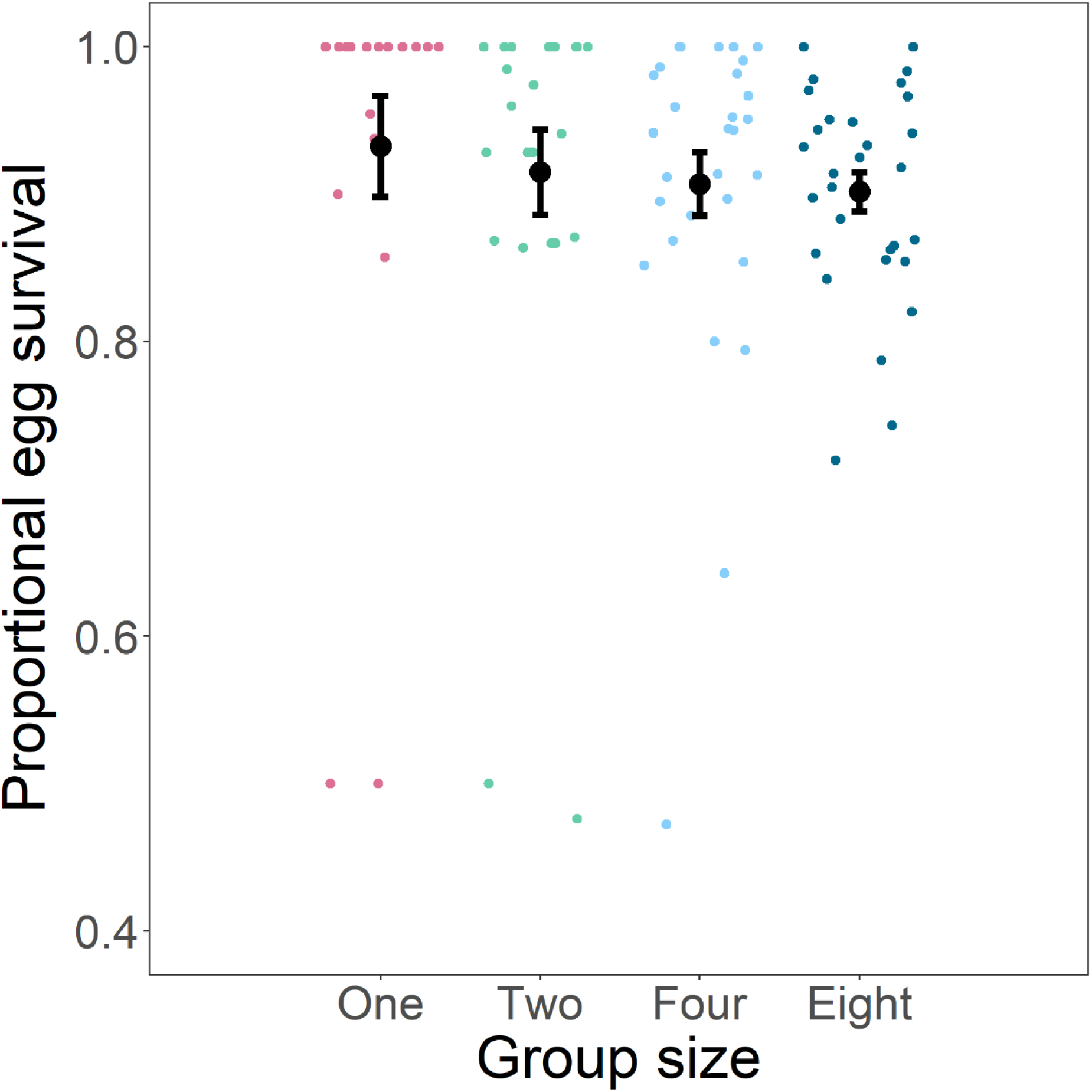
No evidence of fitness effects of social environment. There was no effect of social group size on egg-adult viability (solitary: N = 20 vials; paired: N = 24 vials; groups of 4: N = 29 vials; groups of 8: N = 30 vials). Means (large dot) and standard errors are shown in black.

**Table S1.** Edge effect egg laying location decisions were not overridden by adult social group size or existing egg locations. Effect of social density and previous egg laying patterns on female fecundity and egg clustering. The proportion of vials where eggs laid were observed to be clustered with the existing egg (main bodies in direct contact) and laid at the edge of the vial (within 0.5cm from the vial circumference).

|  |  | Proportion of eggs laid<br>at the edge (%) | Proportion of eggs<br>clustered (%) |
| --- | --- | --- | --- |
| Social treatment | Egg location |  |  |
| Solitary females | None (N = 33 vials) | 100 | - |
|  | Edge (N = 22 vials) | 100 | 0 |
|  | Centre (N = 15 vials) | 100 | 0 |
| Groups of four | None (N = 17 vials) | 94.1 | - |
|  | Edge (N = 17 vials) | 93.8 | 0 |
|  | Centre (N = 13 vials) | 100 | 0 |

**Table S2.** Egg laying patterns of females were significantly non-random. Comparison of the clustering proportion model outputs and empirical egg laying patterns observed in *D. melanogaster*. Shown are Kolmogorov-Smirnov P value outputs for the comparison of empirical data to expected clustering preference values (К = 0 to К = 1, in 0.1 increments) rounded to two decimal places. Values that show the most likely clustering preference are highlighted in bold.

| Social treatment | K values |  |  |  |  |  |  |  |  |  |  |
| --- | --- | --- | --- | --- | --- | --- | --- | --- | --- | --- | --- |
|  | 0.0 | 0.1 | 0.2 | 0.3 | 0.4 | 0.5 | 0.6 | 0.7 | 0.8 | 0.9 | 1.0 |
| Solitary | 0.09 | 0.34 | 0.44 | **0.48** | 0.46 | 0.29 | 0.17 | 0.06 | 0.02 | 0.03 | 0.01 |
| N = 20 vials |  |  |  |  |  |  |  |  |  |  |  |
| Paired | 0.01 | 0.20 | 0.34 | 0.43 | **0.43** | 0.38 | 0.22 | 0.15 | 0.07 | 0.04 | 0.01 |
| N = 25 vials |  |  |  |  |  |  |  |  |  |  |  |
| Groups of four | 0.00 | 0.08 | 0.16 | 0.25 | **0.29** | 0.29 | 0.23 | 0.13 | 0.05 | 0.02 | 0.00 |
| N = 29 vials |  |  |  |  |  |  |  |  |  |  |  |
| Groups of eight | 0.00 | 0.00 | 0.07 | 0.15 | 0.20 | **0.21** | 0.15 | 0.08 | 0.03 | 0.01 | 0.00 |
| N = 30 vials |  |  |  |  |  |  |  |  |  |  |  |

